# Embryonic and larval development of the Pacific saury *Cololabis saira*: Distinctive characteristics of a rapidly growing beloniform fish

**DOI:** 10.64898/2026.02.10.705229

**Authors:** Rie Kusakabe, Shinya Yamauchi, Shigehiro Kuraku

## Abstract

**Background:** Pacific saury *Cololabis saira* is one of the important food resources drawing attention for its recent rapid decline of catch. Their life cycle and embryonic development have been largely unknown. It is important to clarify how the habitat and reproduction of this species have been affected by the global changes of aquatic environment.

**Results:** We obtained fertilized eggs of *C. saira*, by spontaneous spawning and artificial fertilization, and observed the embryonic development up to larval stages. Embryonic stages are documented with major periods of developmental events; cleavage, gastrulation (epiboly) and somitogenesis and organogenesis. Remarkably, segmentation of somites starts in the middle of epiboly, unlike other well-documented teleost species such as zebrafish and medaka. Morphological changes in larval stage up to feeding juvenile is also described. Growth speed of larval Pacific saury is dramatically rapid, in comparison to closely related beloniform fish such as medaka.

**Conclusions:** In comparison to medaka, early embryogenesis of saury proceeds slowly, although being followed by early onset of somitogenesis. This might be partly responsible for the rapid growth into adult (larger than 20 cm in body length) in only half a year. Further studies on embryonic development will uncover the molecular mechanisms underlying the characteristics of Pacific saury as an excellent source of nutrition and as an indicator of major environmental changes such as global warming.

## 1. Introduction

Sauries are marine sword-shaped pelagic fish living in tropical and temperate waters. In east Asia, sauries inhabit North Pacific Ocean, the Sea of Japan and the Sea of Okhotsk. Pacific saury *Cololabis saira* had been one of the common and inexpensive marine food sources served in daily cuisines in Japan ^1^. In the last decades, however, catch of *C. saira* has been declining dramatically, leading to a sharp rise of the market price. This might be partly due to the recent global environmental changes associated with elevated temperature of sea water. Pacific saury stays in the northern Pacific during the summer and starts to migrate toward the south in early fall (Fuji et al., 2019, NPFC). Spawning of sauries peaks during winter and lasts into early summer next year. Larval sauries grow rapidly into adults and become sexually mature in about 240 days after hatching ^2^. During this period, adult sauries grow up to 25 to 35 cm in body length.

Sauries belong to the order Beloniformes that exhibits particularly remarkable characteristics with respect to the morphological diversity as well as to the history in fisheries industry. Beloniformes comprises of 298 species including medakas, needlefish, flying fishes, halfbeaks and sauries, associated with a variety of highly functionalized body shape. For example, flying fish have wing-like pectoral fins that provide gliding ability in the air. Their caudal fin with longer ventral edge assists the body to take off from the water. Halfbeaks show large difference in the length of jaws; their lower jaw is extremely longer than the upper jaw. Although phylogenetically nested within the family Belonidae ^3^ and NCBI Taxonomy), the Pacific saury is traditionally classified in the family Scomberesocidae in morphological taxonomy due to the presence of dorsal and anal finlets ^4,5^. Dorsal and ventral finlets are also common to scombrid fish and are thought to contribute to the high swimming speed ^6^.

With respect to the developmental processes, only a few teleost species have been studied in detail. The developmental processes of teleosts have been stereotyped by that of zebrafish *Danio rerio*, a particularly prevailing non-mammalian developmental model ^7^. Its rapid embryogenesis and transparency of embryonic body have allowed researchers to identify the peculiar mode of gastrulation, the epiboly, followed by tailbud formation, somitogenesis and organogenesis. However, zebrafish belongs to Ostariophysi which is included in Otocephala, a cohort that diverged 230 mya from the rest of the teleosts including Euteleostei (Figure 1A). Only a few literatures for description of developmental processes of Euteleosts can be found, including that of medaka *Oryzias latipes* ^8^, rainbow trout ^9^ and the false crownfish^10^. These studies have implied potential deviation, either subtle or obvious, from the established scheme of teleost development based mainly on zebrafish. Further studies covering wider range of teleost orders would be necessary to clarify the key developmental features underlying the evolutionary success in a particular order of euteleosts.

**Figure 1.**
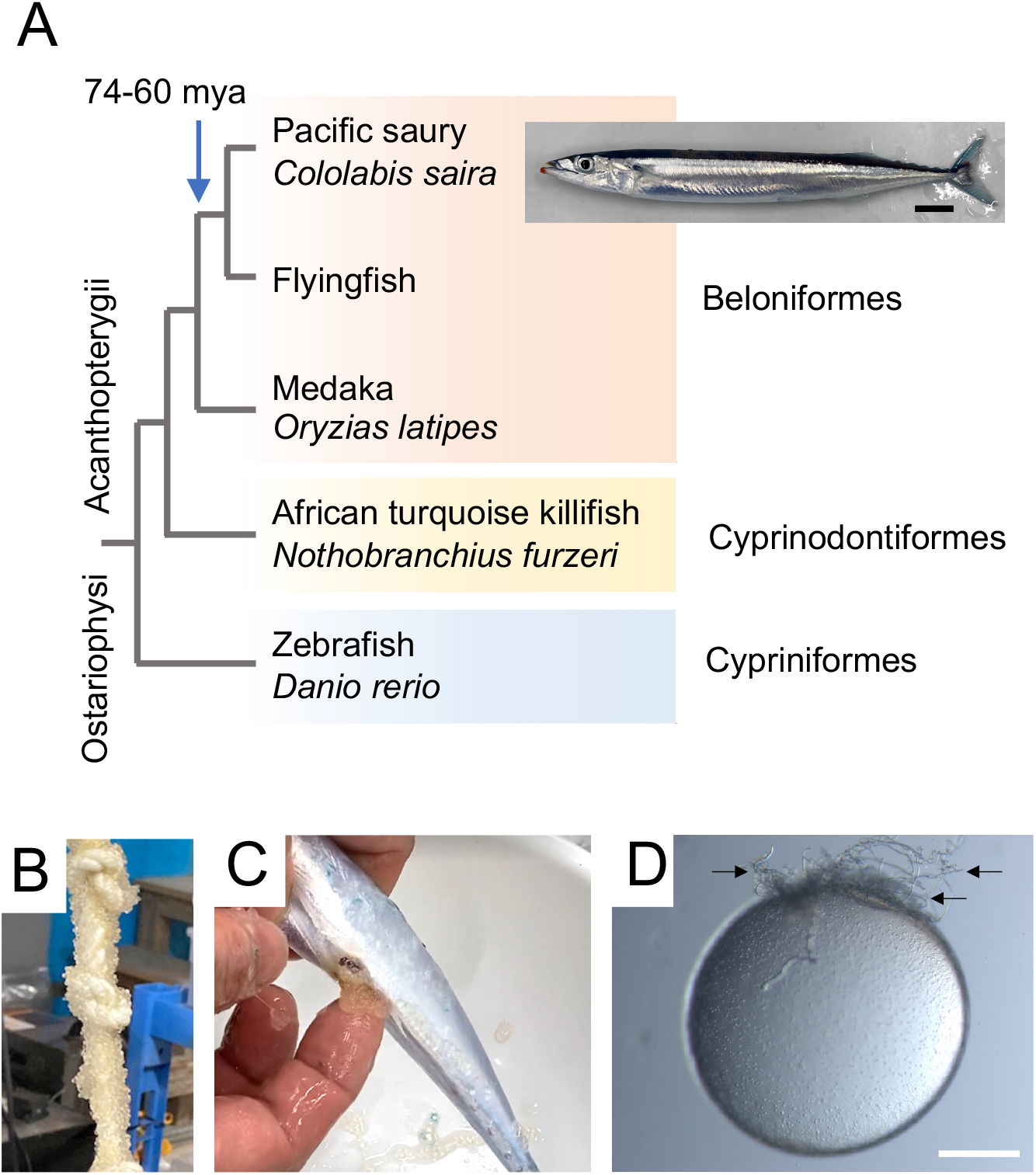
*C. saira* and its eggs obtained in the aquarium. A. Phylogenetic relationships of Belonoid and major teleost model species (medaka, killifish, zebrafish). The pacific saury and the Japanese medaka diverged 74 to 60 million years ago (mya). Scientific names of representative species are provided in italics. Scale bar: 2cm. B. Eggs naturally spawned on the surface of a rope hung in the exhibition tank. C. Eggs deposited from a female by manual pressure. D, An unfertilized egg of *C. saira*. Arrows indicate the clipped attaching filaments at the animal pole. Scale bar: 0.5mm.

Fertilized eggs of *C. saira* can be found tangled with drifting algae. After collection, these eggs can be reared in the laboratories ^2^. A series of persistent efforts led to the successful maintenance and maturation of the hatched larvae in several research institutes in Japan ^2^ and references therein). Growth process of post-hatch *C. saira* larvae has been documented based on the observation of individuals hatching in the fish tank ^2^. When placed in appropriate water temperature and fed to satiation, juvenile sauries grow into sexually mature adults that are around 20 cm in body length within one year after fertilization (personal communication). This represents one of the peculiar characteristics of the Belonoid fish, the varied speed of growth; sauries grow remarkably fast, whereas adult medakas are only about 3 cm in adult body length. It is also notable that both sauries and medaka exhibit high swimming capabilities upon hatching and start seeking food immediately, suggesting that their embryonic development achieves fully mature locomotive and digestive functions.

Driven by the growing concern about the pelagic fish as a critical part of marine resource, there has been a demand for characterization of life cycles and genomic features of yet unexplored Belonoid species. Recently whole-genome sequence of *C. saira* has been analyzed in detail by three independent research groups, characterizing genomic properties of this species^11-13^. However, morphogenetic processes in early development of *C. saira* have yet to be clarified. In order to further examine morphogenetic characteristics of sauries, a standardized developmental stage table would be necessary.

The present study is the first report of developmental table of a scomberesocid from the first cleavage to the feeding larval stage. Fertilized eggs are obtained either by spontaneous spawning in the aquarium tank during the breeding season (Figure 1B) or by artificial fertilization (Figure 1C and D). Overall morphology of *C. saira* embryos resembles that of medaka, although the relative timings of morphogenetic events show difference between the two species. We discuss the differences in developmental events and growth speed between *C. saira* and *O. latipes*, the two beloniform species that diverged around 74-60 million years ago (Figure 1A)^3,11,14,15^.

## 2. Results

### 2.1 Fertilization and early cleavage of *C. saira*

Spawned unfertilized eggs of *C. saira* are about 1.8mm in diameter, slightly larger than medaka *O. latipes* eggs (Figure 1D). A *C. saira* egg is enclosed in a thick transparent chorion (egg envelope) associated with long attaching filaments (Figure 1D). These filaments are located at the future animal pole, unlike medaka eggs that have attaching filaments at the vegetal pole. Unlike that of medaka eggs, which have hundreds of short non-attaching filaments all over the egg surface ^8^, the chorion surface of *C. saira* is smooth except the part where the attaching filaments are located.

Small oil particles are scattered evenly in the yolk (Figure 1D). In the aquarium tank, eggs are spawned on the substrate (spawning beds) (Figure 1B) and immediately inseminated by males. The spawning behavior starts in the morning, as soon as the lights turn on, and lasts for several hours, the spawning bed becomes covered with clutches of fertilized eggs (Figure 1D). In this study, these naturally spawned eggs were served for observation of blastula and later stages described below. Earlier stages, i.e., cleavage, was documented based on the observation of artificially fertilized eggs.

Newly fertilized eggs have an ovoid shape, with the animal-vegetal axis slightly longer (Table 1 and Figure 2A). After fertilization, oil particles are no longer visible on the egg surface. Unlike in many of other teleost species such as medaka and zebrafish, fertilization in the saury triggers a very slight, if any, elevation of chorion (which is now called the fertilization envelope); thus, there is no apparent space between the egg cytoplasm and surrounding chorion (Figure 2A). 30 minutes after fertilization, non-yolk cytoplasm appears at the animal pole and gradually increase its height. The cytoplasm undergoes the typical discoidal cleavage. At 2 hours post-fertilization (hpf), the first cleavage furrow divides the cytoplasm equally (Figure 2B). The furrow divides the blastodisc, but not the yolk of the egg, and thus incomplete (meroblastic). Cleavages continue synchronously every 30 to 50 minutes at 16°C to produce 4, 8, 16, 32 cells (Figure 2C-F and Table 1). Up to 64-cell stage, cells are organized in a single layer connected to the yolk (Figure 2G). Further cleavages lead to the stratification of the cells that form a multi-layered blastodisc (Figure 2H). This mass of cells becomes surrounded by a ring of cytoplasm, the germ ring (Figure 2I). In the following epiboly movement, the germ ring serves as a leading edge of the expanding blastodisc.

**Table 1.** Stages of embryonic development of *C. saira*. (at 16°C)

| Stage | Figure | Time | Description | # of somites |
| --- | --- | --- | --- | --- |
| 1-cell | 2A | 1-2 hpf | First cell appears at the animal pole |  |
| 2-cell | 2B | 2 hpf | 2 blastomeres |  |
| 4-cell | 2C | 3 hpf | 2 x 2 blastomeres |  |
| 8-cell | 2D | 4 hpf | 2 x 4 blastomeres |  |
| 16-cell | 2E | 4.5 hpf | 4 x 4 blastomeres |  |
| 32-cell | 2F | 5.3 hpf | Cell orientation not clear |  |
| 64-cell | 2G | 6 hpf | Cells form a single-layered blastodisc |  |
| Early blastula | 2H | 17 hpf | Cells form a multi-layered blastodisc |  |
| Late blastula | 2I | 19 hpf | Germ ring encircles the blastodisc |  |
| Germ ring | 3A | 28 hpf | Germ ring becomes wide and thick at one side |  |
| Shield | 3B & 3C | 39 hpf | Embryonic shield appears |  |
| 30% epiboly | 3D | 42 hpf | Embryonic axis appears |  |
| 50% epiboly | 3E & 4A | 45 hpf | Primordial eyes are formed at the rostral end.<br>Somite segmentaion starts | 3 |
| 60% epiboly | 3F | 48 hpf | Kupffer's vesicle appears<br>At least 3 somites are visible |  |
| 70-80% epiboly | 3G & 4B | 54 hpf | Kupffer's vesicle becomes larger | 8 |
| 90% epiboly | 3H | 60 hpf | Blastopore is about to close and the yolk is completely internalized<br>Germ ring is still visible |  |
| 100% epiboly | 4C | 70 hpf | Blastopore is completely closed | 12 |
| Pre-tailbud | 5A & 6A | 3 dpf | Mid-hind brain boundary is visible. Otic vesicles appear. Heart starts to beat at the ventral side of the head. Blood cells (with no color) are visible being pumped in and out of the heart. | 35 |
| Early tailbud | 5B, 6B & 7A | 3 dpf | Tail bud appears and elongates. 2 otoliths (ear stone) appear in each of the otic vesicle. Pectoral fin buds start to develop. | 57 |
| Mid tailbud | 5C | 4 dpf | Tailbud overlaps with heart. |  |
| Late tailbud | 5D | 5 dpf | Tail overlap with pectoral fin. Pigmentation starts on the epidermis of the yolk surface. Pectoral fins move frequently. |  |
|  | 6C | 6 dpf | Heart is shaped in a complete tube-like structure. Red blood cells are found in the blood stream. |  |
|  | 5E & 6D | 7 dpf | Pale black pigmentation covers the head, eyes and dorsal side of the trunk/tail. Mouth opens. |  |
| Hatching | 6A & 6B | 9-11 dpf | Hatching starts. |  |
| Early larva | 6C | 12 dpf | All the embryos hatch by this time. Lower jaw elongates and becomes protrusive. |  |
|  | 6D | 14 dpf | Abdominal wall completely covers the yolk. |  |
|  | 6E | 25 dpf | Larvae can survive without feeding up to this stage. |  |

**Figure 2.**
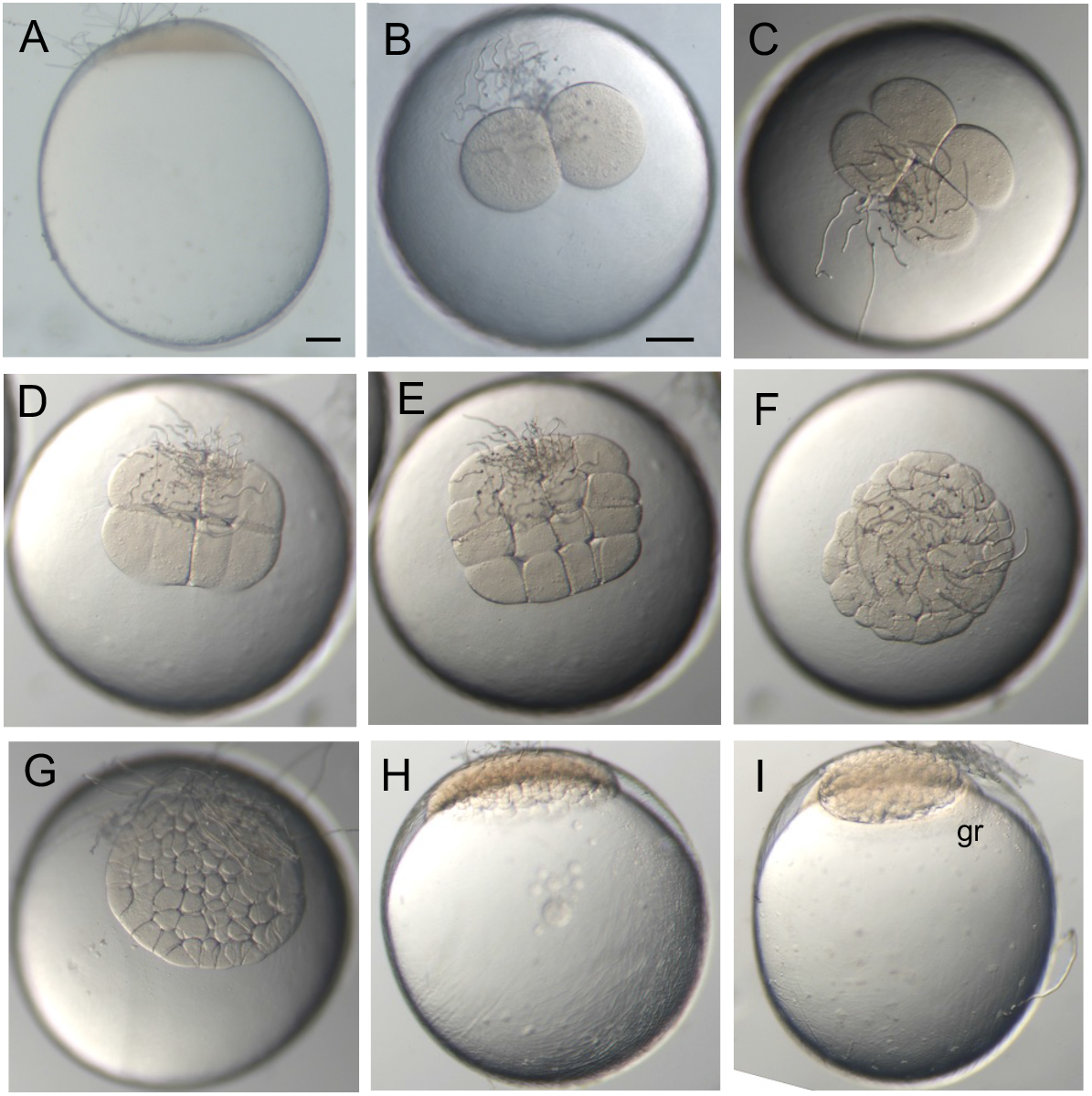
Early embryogenesis of *C. saira*. A. 1-cell stage, animal pole to the top. B. 2-cell, C. 4-cell, D, 8-cell and E. 16-cell stages viewed from animal pole. F and G, single-layered blastodisc viewed from animal pole. H. multi-layered blastodisc, animal pole to the top. I. germ ring (gr) appears at the edge of the blastodisc. gr, germ ring. Scale bars: 0.2mm.

### 2.2 Epiboly stage and formation of embryonic body

At 24 hpf (hours post-fertilization), the blastodisc starts to expand toward vegetal pole (epiboly; Figure 3). At the onset of epiboly, one side of the germ ring becomes widened, marking the future dorsal side of the embryo (Figure 3A). When blastodisc covers the animal portion of the yolk (Figure 3B), the dorsal germ ring cells intercalate and form a triangle-shaped thickening, the embryonic shield (Figure 3C)^16^.

**Figure 3.**
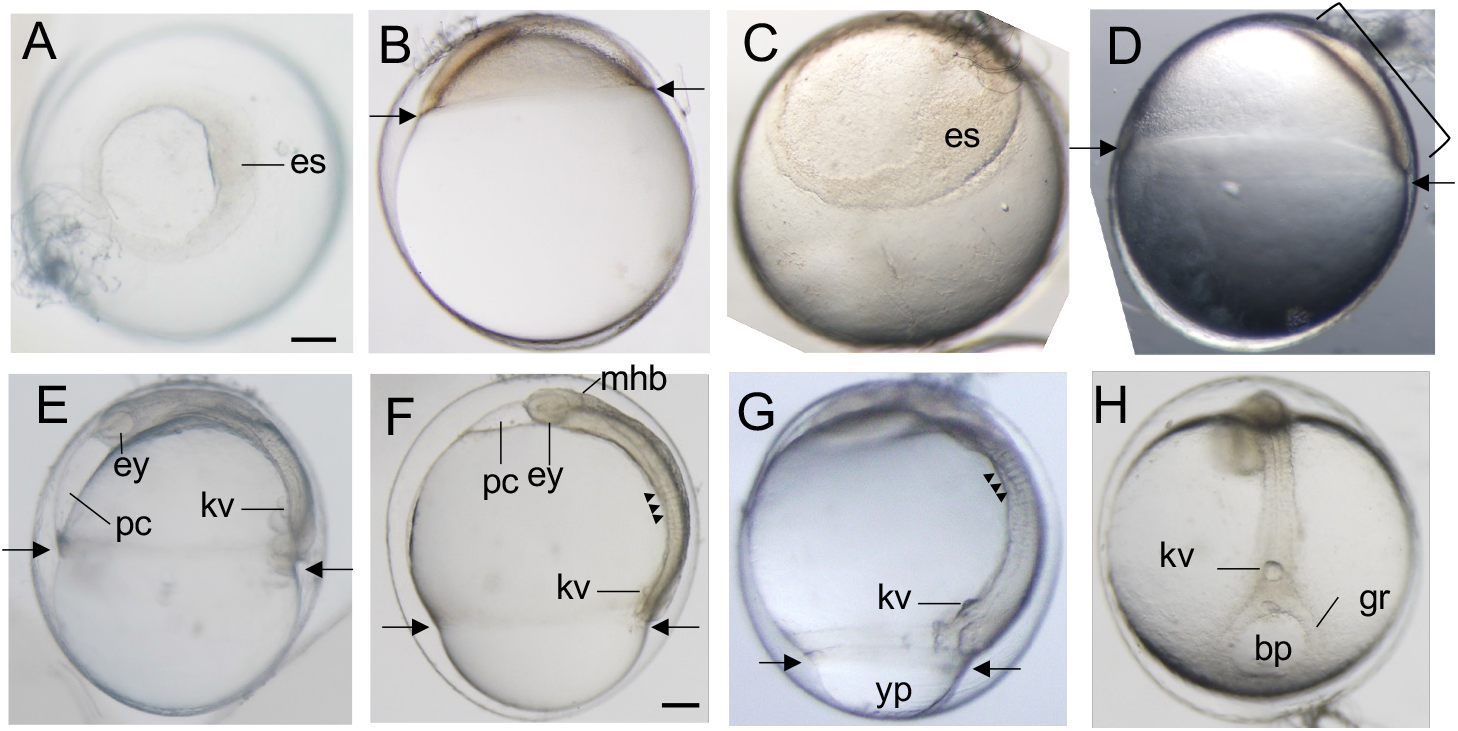
Epiboly (gastrulation) of *C. saira* In all figures, arrows indicate the level of extending edge of the blastoderm. A. At the onset of epiboly, one side of the germ ring becomes thick and wide, marking future embryonic shield (es). B and C. Shield stage. B. Right lateral view. C. Tilted animal view of the blastoderm. Embryonic shield (es) is clearly visible. D-H, Progression of epiboly. Arrows indicate the level of extending edge of the blastoderm. D, 30% epiboly. Left-lateral view. Bracket indicates the forming embryonic body. E. 50% epiboly. Left lateral view. Eyes (ey) are forming anteriorly to the body axis. F. 60% epiboly. Forming somites (indicated by triangles) and Kupffer’s vesicle (kv) are visible. G. 80% epiboly stage. The caudal part of yolk plug (yp) bulges out of the blastopore. H. A caudal view of 95% epiboly. Blastopore (bp) is now closing. es, embryonic shield; ey, eye; kv, gr, germ ring; Kupffer’s vesicle; mhb, mid-hindbrain boundary; pc, pericardial cavity; yp, yolk plug. Scale bar; 0.2mm.

The embryonic shield continues to extend anteriorly toward the center of the blastodisc as the cells in the lateral blastodisc and germ ring are added to form the axis of embryonic body (Figure 3D). When the extending blastodisc covers 50% of the yolk, the embryonic body is clearly visible with the anterior head associated with rudimentary eyes (Figure 3E). At 70% epiboly (Figure 3F), the Kupffer’s vesicle forms, a potential early sign of posterior growth and left-right patterning ^17,18^. Kupffer’s vesicle initially appears as a cluster of multiple small vesicles, and the fuse to a single large vesicle. As the epiboly further proceeds, the Kupffer’s vesicle becomes larger and placed at the ventral side of the future tailbud (Figure 3G and H). At later stages of the epiboly, the germ ring at the edge of the blastoderm exhibits robust contractile activity to close the blastopore at the vegetal pole. In some individuals, the germ ring pinches the inner yolk mass so that it transiently bulges out to form a yolk plug (Figure 3G). When epiboly finishes, the yolk is completely internalized and the germ ring further contracts to close the blastopore (Figure 3H).

### 2.3 Somitogenesis, tail elongation and onset of pigmentation

Somite segmentation starts at 50% to 70% epiboly stage (Figure 3F and 4A). At this stage, eyes and the forming brain are visible. Although the precise timing varies between individuals, the onset of somitogenesis in *C. saira* coincides with the ongoing epiboly, unlike that in zebrafish and medaka^7,8,10^. At 50% epiboly, 2 or 3 somites were recognizable. Somitogenesis proceeds in anterior-to-posterior direction in concert with progression of epiboly (Figure 4B and B’). Somite number increases by roughly one somite per 2 hours. When the blastopore closes, the embryo has more than 12 somites at each lateral side of the body (Figure 4C and C’). During the tail elongation (described below), somite formation accelerates to produce one somite per 30 minutes.

**Figure 4.**
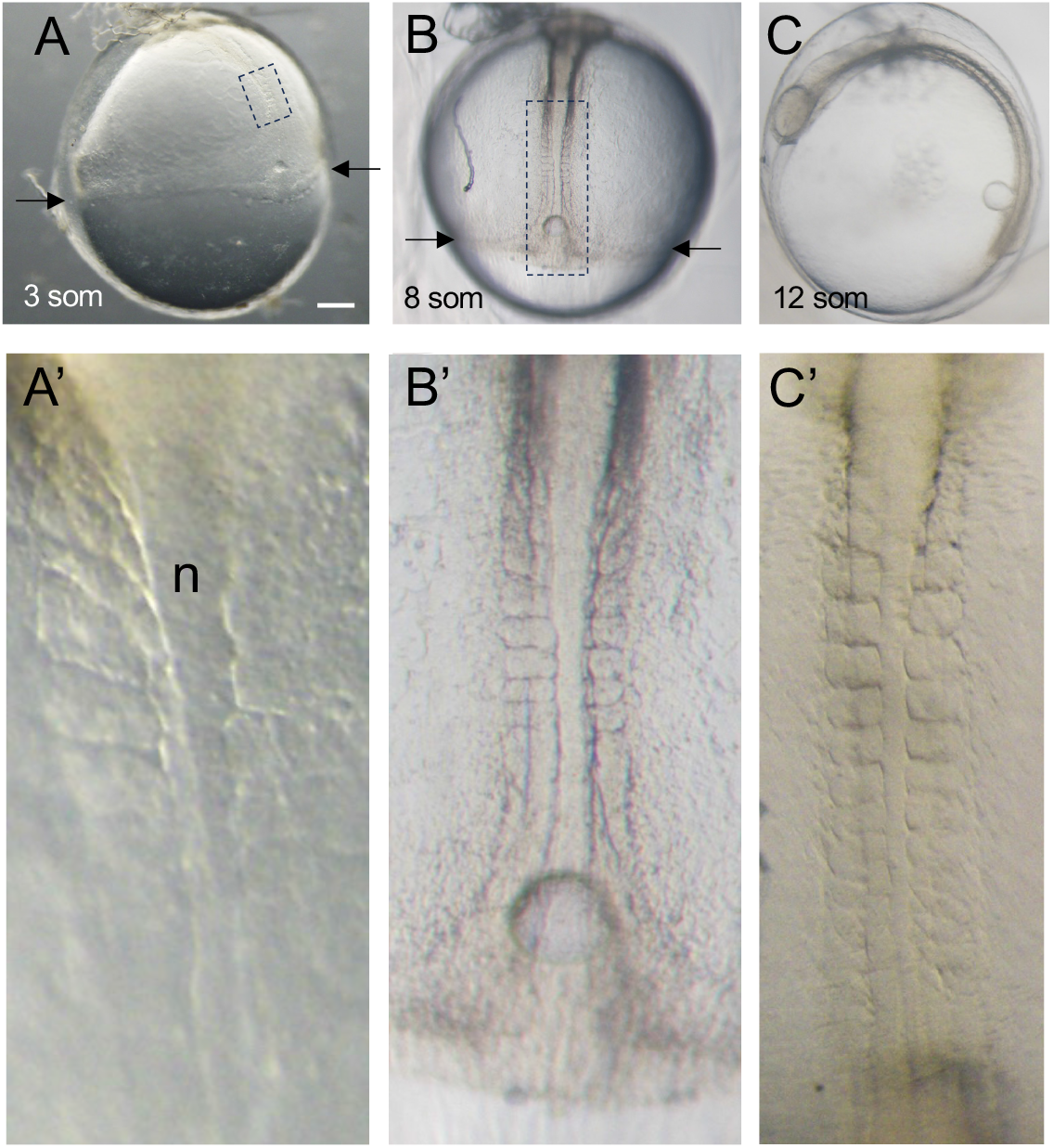
Somitogenesis of *C. saira* A. An embryo at 50% epiboly associated with 3 pairs of somites visible. B. Dorsal view of an embryo at 70% epiboly with 8 pairs of somites. A’ and B’, The magnified dorsal views of areas framed with dotted line in A and B, respectively. Anterior is to the top. C. An embryo at 100% epiboly has about 12 pairs of somites. C’. The same embryo in C viewed from dorsal side. Anterior is to the top. n, notochord. Scale bar; 0.2mm.

When the epiboly is completed, the head is associated with prominent eyes, midbrain, hindbrain and mid-hindbrain boundary (Figure 5A). Shortly afterwards, tailbud structure becomes obvious (Figure 5B). Inside the egg chorion, the tail elongates posteriorly toward the head and, on 4 days post-fertilization (dpf), reaches the heart that has already started pumping (Figure 5C). In the late tailbud stage, the tip of the tail passes through either right or left side of the head and reaches the forming pectoral fins (Figure 5D). At this stage, black pigmentation is observed scattered on the surface epidermis of the yolk and head (arrows in Figure 5D). Unlike medaka with retinal pigmentation starts well ahead of the body pigmentation ^8^, the Pacific saury has unpigmented eyes until late tailbud stage (Figure 5C and D).

**Figure 5.**
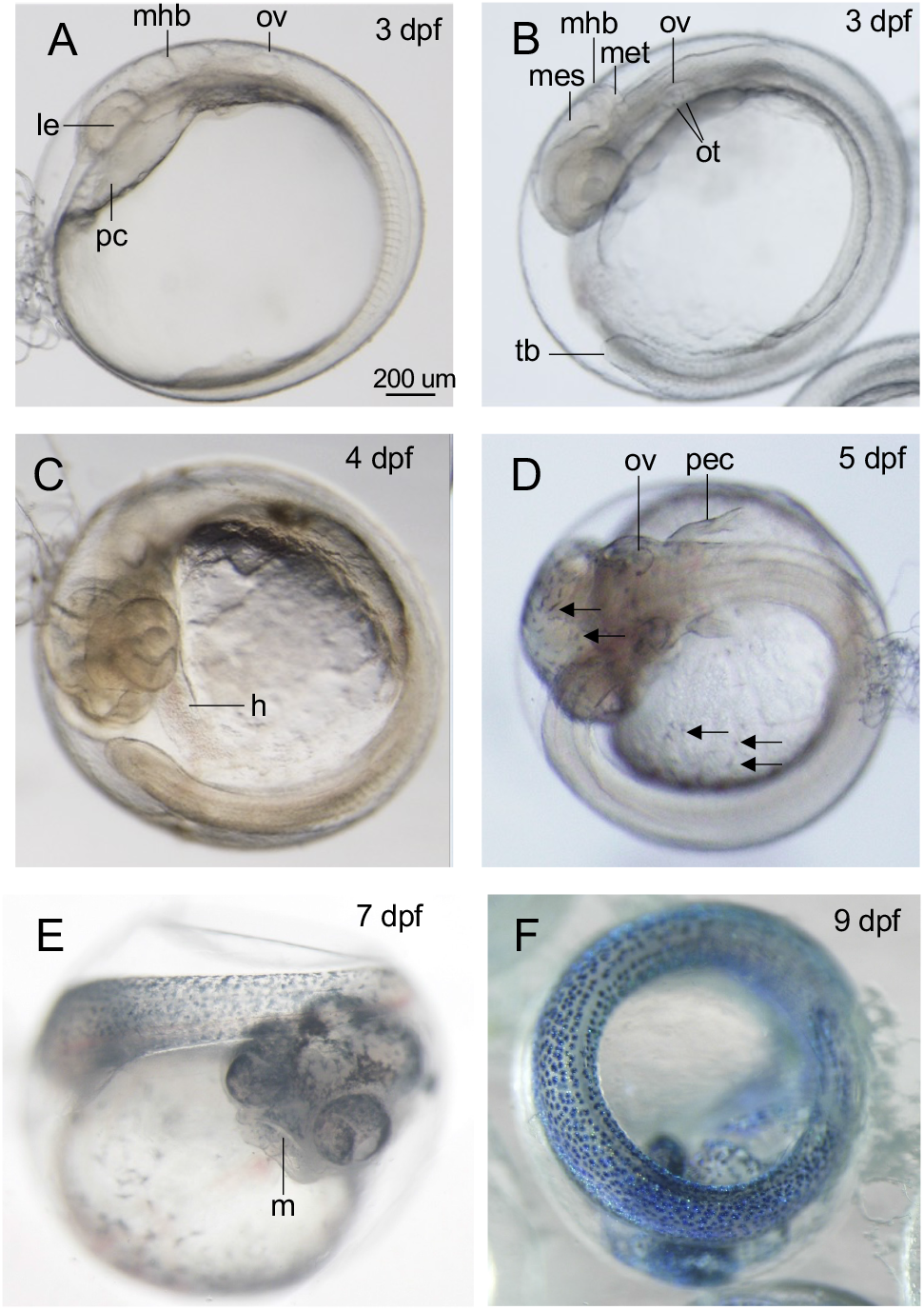
Tailbud elongation and later development A-C. Tail elongation stage embryos viewed from the left side. A. At early 3 dpf, otic vesicle (ov) forms anteriorly to the somites. le, lens. B. At later 3 dpf, Tailbud (tb) is clear. The otic vesicle now has 2 otoliths (ot). C. 4 dpf. Tailbud elongates and now overlaps with the heart primordium (h). D. the tail further elongates and wraps around the yolk. The tip of the tail now reaches the forming pectoral fins (pec). Arrows indicate black pigment cells. E. At 7 dpf, mouth (m) opens. F. At 9 dpf, near-hatching embryo with numerous blue pigmentation in the tail. h, heart primordium; m, mouth; mhb, mid-hindbrain boundary; mes, mesencephalon; met, metencephalon; ov, otic vesicle; ot, otolith; pc, pericardial cavity.

As the tail elongates further and passes the pectoral fins, robust pigmentation starts in the head, eyes and tail (Figure 5E). The initial pigment color appears entirely black, probably due to the melanophore development. In hatching period, the embryos or early larvae exhibit characteristic blue color (Figure 5F). We have also analyzed the physiological nature of the pigment cells of the larvae, but this will be reported elsewhere.

### 2.4 Organogenesis

#### Development of the head

Head structure becomes evident by the appearance of bilateral optic vesicles (‘ey’ in Figure 3E) at 50% epiboly. Thus the appearance of the rudimentary eyes occurs much earlier in the saury than in the medaka in which primordial eyes appear after the completion of epiboly ^8^. By the end of epiboly, but before the appearance of tailbud, lens develops in the center of the eye (Figure 5A). Pigmentation of the eyes starts at around 7 dpf simultaneously with the increase of pigment cells on the head epidermis (Figure 5E).

Another signature of head development is the appearance of brain compartments. Mid-hindbrain boundary (MHB) is visible posteriorly to the forming eyes as early as at 60% epiboly (Figure 3F). In the tailbud stage, MHB clearly marks the division between mesencephalon and metencephalon (Figure 5A and B). Initial otic vesicles are formed posteriorly to the MHB (Figure 5A). When tailbud appears at late 3 dpf, two otoliths (ear stones) appear in the ventromedial aspect of each otic vesicle (Figure 5B and 7A). At around 7 dpf, mouth opens at the rostral end of the head and movement of the lower jaw is observed (Figure 5E).

#### Cardiovascular development

At 50% epiboly, the heart formation starts with the presence of the pericardial cavity located anterior to the head (Figure 3E and F). It is a large space that will enclose the future heart. At the end of epiboly stage, the pericardial cavity moves posteriorly and becomes internalized underneath the head (Figure 5A). At 3 dpf, primitive heart region is recognizable with the constant pulsation (Figure 6A). At 4 dpf, the heart forms a single tube with the blood containing globular blood cells flowing into it through common cardinal veins (ccv) (Figure 6B). These veins form two branches extending over the right and left sides of the yolk. This initial blood stream has no color and gradually turns red as the red blood cells differentiate (Figure 6C). At 6 dpf, the heart is now composed of two chambers, the atrium and the ventricle, pumping alternately (Figure 6C). The yolk surface becomes covered by the network of blood vessels that carry the blood stream in posterior-to-anterior direction (Figure 6D). In the trunk, the dorsal aorta lies on the midline underneath the notochord and extends into the tail (Figure 6E). In the tail, the dorsal aorta is defined as the caudal aorta along which the caudal vein runs as the return route toward the heart (Figure 6E’). The caudal vein is connected to marginal vein that lies underneath the digestive tract (Figure 6E).

**Figure 6.**
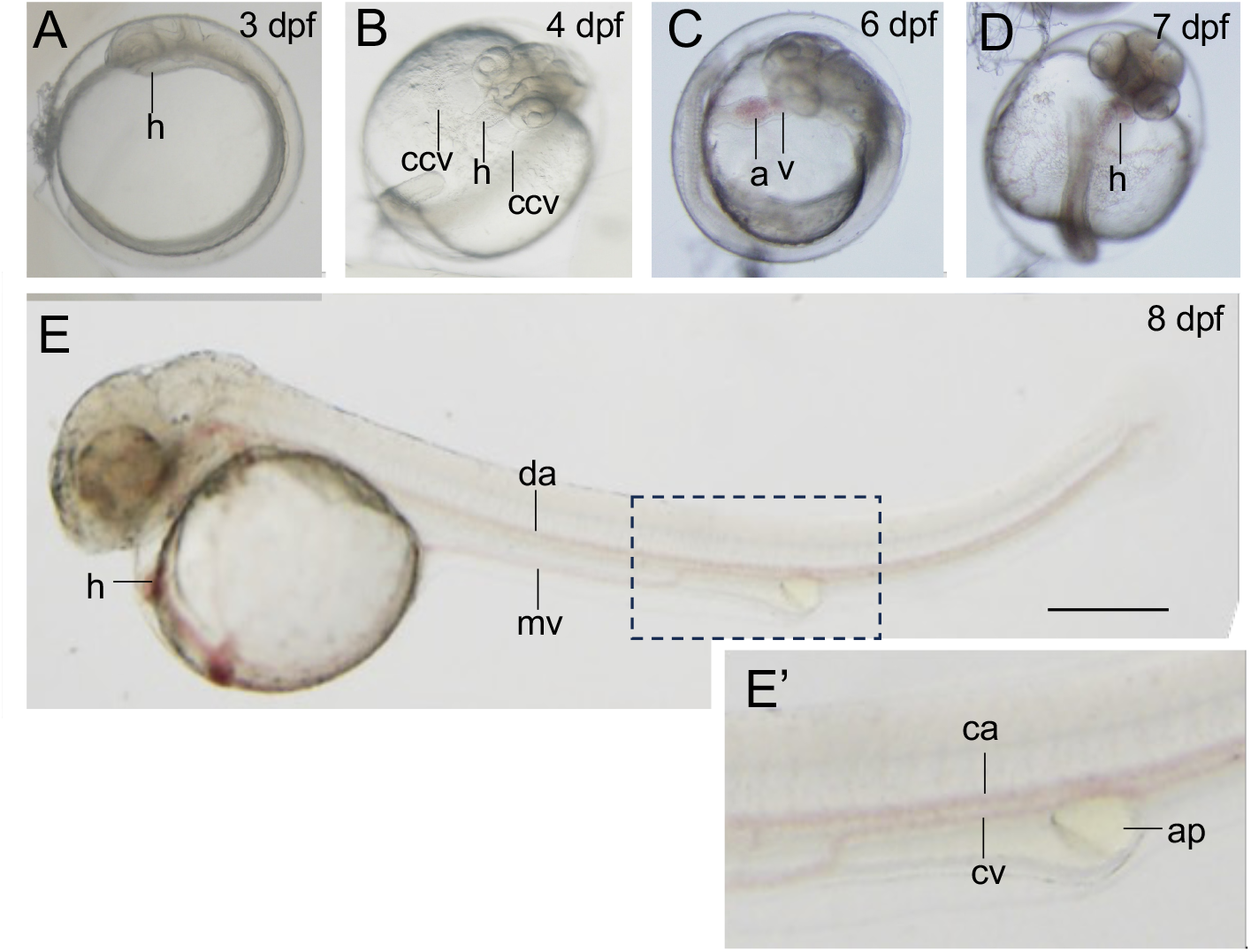
Cardiovascular development A. At 3 dpf, primordial heart starts to beat. B. At 4 dpf, the heart is now tube-shaped structure connecting to common cardinal veins (ccv). C. 6 dpf. Blood stream now contains red blood cells. D. At 7 dpf, the surface of the yolk is extensively vascularized. E. An 8 dpf embryo manually dechorionated. A single medial dorsal aorta (da) extends beneath the notochord. E’, magnified view of the dashed frame in E. The dorsal aorta caudal to the anal pore is called caudal artery (ca). Ventrally to the caudal artery, caudal vein (cv) forms and drains into the marginal vein (mv). a, atrium; v, ventricle; ap, anal pore; ca, caudal artery; ccv, common cardinal vein; cv, caudal vein; da, dorsal aorta; h, heart; mv, marginal vein.

#### Pectoral fins

In early tailbud stage, the pectoral fin development is observed bilaterally in parallel to the body axis at the level of second and third somites (Figure 7A). Earliest phase of pectoral fin bud is marked by the appearance of the apical fold, an equivalent tissue to the apical ectodermal ridge in amniotes, at the level of somite 1-3 (Figure 7A)^19^. Pectoral fin buds bulge and enlarge in posterior-to-anterior direction and form fan-shaped structure with the endochondral discs at the center (Figure 7B-D). At around 8 dpf, the pectoral fins are now associated with actinotrichia (pre-fin ray fibers) and exhibit frequent movement even before hatching (Figure 7E). After hatching, pectoral fins swing regularly as the young larvae swim, probably contributing to the maintenance of posture.

**Figure 7.**
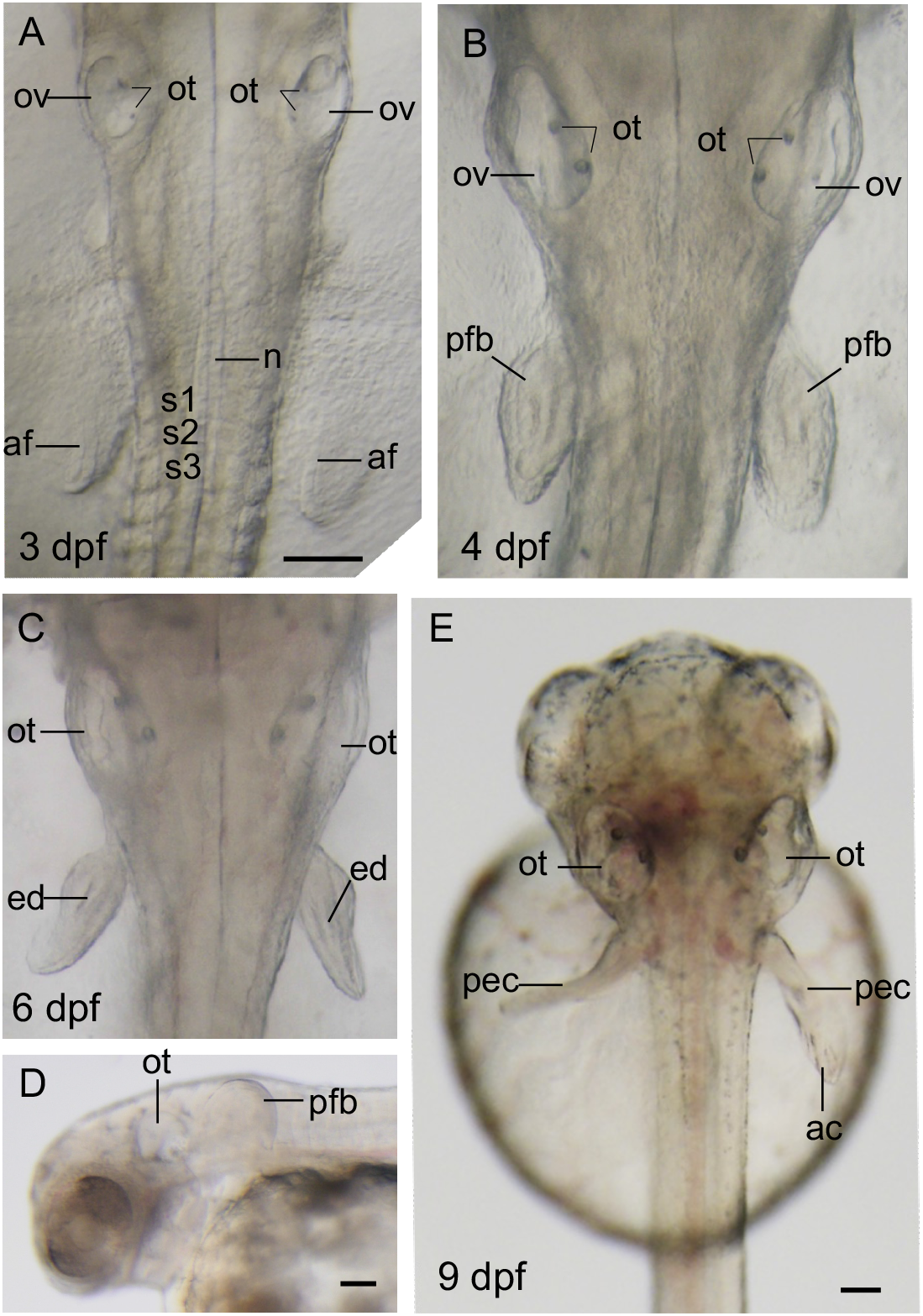
Pectoral fin development A-C, E. Dorsal views of the embryos at otic vesicle to pectoral fin level. A. At 3 dpf, apical fold (af), the future lateral edge of pectoral fin bud, is visible as a swelling of the surface epidermis distally to the lateral body wall. s1 – s3, somites 1 to 3. B. At 4 dpf, pectoral fin buds (pfb) form. C. 6 dpf. Endoskeletal discs (ed) are clearly visible. D. Left-lateral view of a 6 dpf embryo. Pectoral fin bud has grown into a fan-shaped protrusion. E. A 9 dpf embryo immediately after hatching. Pectoral fins (pec) are now associated with actinotrichia (ac). ac, actinotrichia; af, apical fold; ed, endoskeletal disc; n, notochord; ot, otolith; ov, otic vesicle; pec, pectoral fin; pfb, pectoral fin bud; s1-s3, somite 1-3. Scale bars: 0.1 mm.

### 2.5 Hatching and early larval stage

*C. saira* embryos develop into larvae that swim and feed actively soon after hatching, similarly to medaka embryos. Embryos right before hatching exhibit active movement within the egg capsule by wriggling the tail. This would facilitate the breakage of chorion and the tail comes out first. In the laboratory condition, hatching starts around 9 dpf in petri dishes at 16 °C. In the aquarium, embryos cultured in the natural sea water (16-18 degrees) normally hatch at 10 to 13 dpf (data not shown). In some clutches, there was up to two days of delay in hatching. This might be caused by the difference in the concentration of the hatching enzyme released from the egg capsule, which can be different between dishes. There is also a variety in the intensity of body pigmentation in hatchlings; some are associated with scarce and scattered pigment cells and look mostly transparent (Figure 7E), whereas others are entirely covered by densely populated pigment cells (Figure 8A). In the latter case, dorsal and ventral sides of the larval body look blue and silver, respectively.

**Figure 8.**
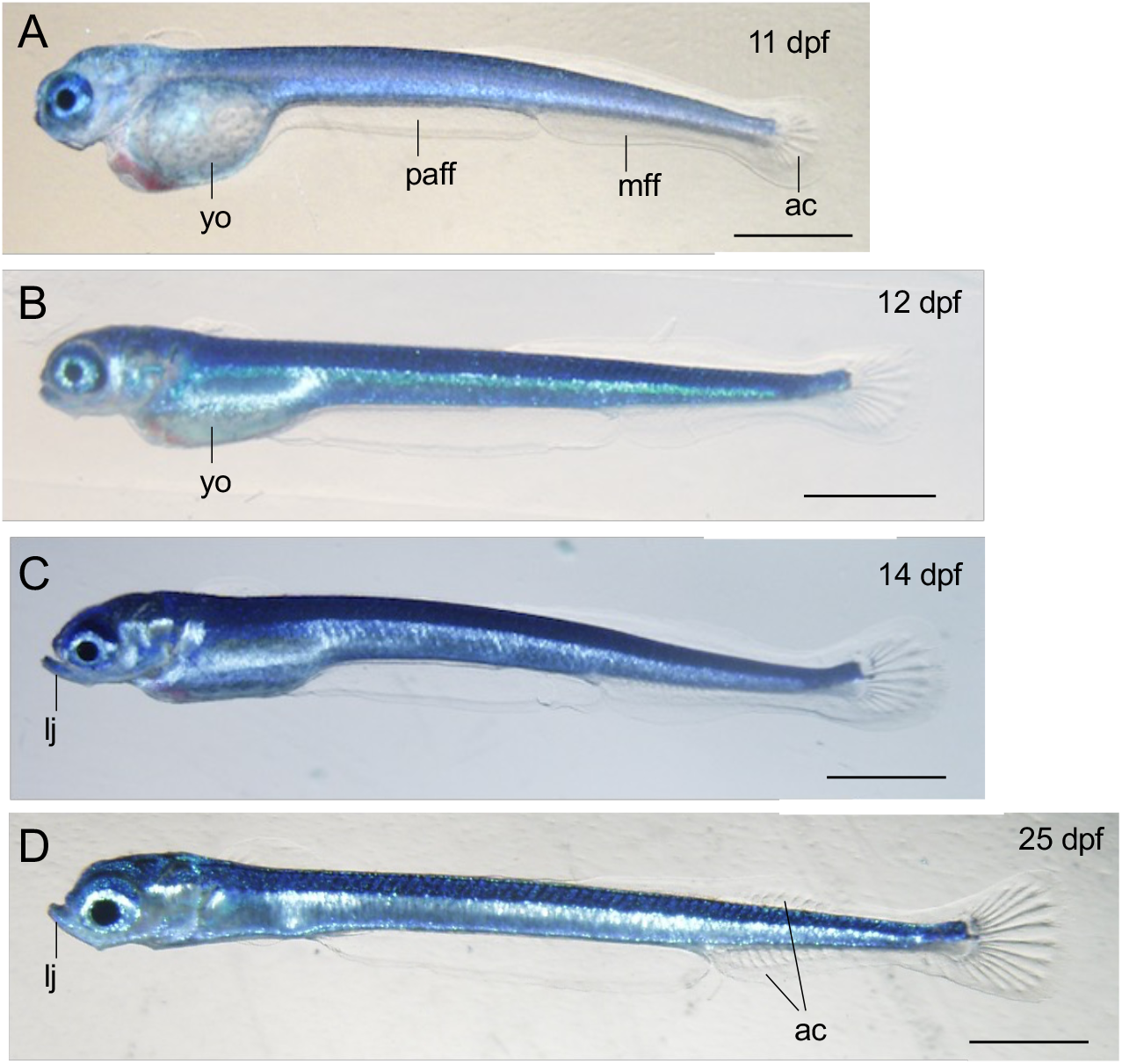
Early larval stages A. A pre-larva before hatching. The chorion was manually removed. Characteristic blue color is now evident. B. At around 12 dpf, hatched larvae sill possess large amount of yolk. C. A larvae one day after hatching. D. A larva 2 weeks after fertilization. The yolk is now completely internalized. ac, actinotrichia; lj, lower jaw; mff, caudal median fin fold; paff, pre-anal fin fold; yo, yolk. Scale bars; A-C, 0.5 mm; D-F, 1 mm.

In the hatchlings, a continuous median fin fold has been formed, consisting of pre-anal and caudal median fin fold (Figure 8A) The former is located underneath at the ventral midline between the caudal end of the yolk sac and the anus. Within the median fin folds, the caudalmost part is the first to develop the actinotrichia (Figure 8A). In later larval stage, actinotrichia also appear in the adjacent anterior regions to the caudal fin at both dorsal and ventral aspects (Figure 8D). Pre-anal fin fold, however, does not form actinotrichia so long as we observed in the laboratory. This is in consistent with the fact that pre-anal fin fold is a transient structure that disappears during late development in actinopterygian species ^20^.

Newly hatched larvae still have a substantial amount of yolk covered by thin abdominal walls (Figure 8B). In the laboratory condition with no feeding, they depend on the nutrition from the yolk and survive up to around 25 dpf (for appropriate feeding condition of larval culture, see Nakaya et al., 2010). Morphology of the mouth starts to exhibit Beloniformes-specific features; the lower jaw extends anteriorly and protrudes over the upper jaw (Figure 8C and D).

## 3. Discussion

In this study, we describe the embryonic and early larval development of the saury *C. saira*, one of the representative marine beloniform species in the North Pacific. Beloniformes include medaka *O. latipes*, which is utilized as a popular model organism of genomics, developmental biology and environmental sciences ^21-24^. To this date, however, early development of no other beloniform species had been reported in detail.

Close examination of *C. saira* embryos enabled us to compare developmental characteristics between sauries and medakas. Eggs of both groups are highly transparent and in similar size range (0.8 – 1.8 mm in diameter) (Kim and Park, 2021). Cleavage pattern (discoidal) and the chronological order of developmental steps (cleavage, blastula, epiboly and tailbud stages) are highly conserved. Hatching of both saury and medaka occurs at around 10 days after fertilization. Hatching larvae have well-developed musculoskeletal system and fins, enabling them to swim actively and seek for food. This feature is supported by the rapid formation of trunk skeletal musculatures, pectoral fins and the mouth ready for food intake. In a clear contrast, embryos of zebrafish, a distantly related Ostariophysi, hatch as early as 48 hpf. Zebrafish hatchlings are premature with respect to the locomotive capability and feeding. At hatching, zebrafish pectoral fins are still at early phase of fin bud formation and thus not mobile. Mouth is already open but still undergoing initial condensation of jaw cartilages, and thus not functional for food intake ^7^. Premature hatching has also been reported for the Midas cichlid fish ^25^. Both zebrafish and cichlids are dependent on yolk as the source of nutrition for an extended period. Thus, relatively late timing of hatching of the saury, as well as medaka, might reflect the early start of feeding that occurs as soon as the larvae are released to the open environment. This is consistent with the previous report that shows the requirement of food abundance for successful maintenance of saury larvae under laboratory conditions ^2^.

Despite the similarities described above, it is remarkable that sauries and medakas grow into adults of considerably different size. Saury larvae grow dramatically fast and become reproductively mature adults of 25 to 35 cm in body length, which takes only 7 to 12 months after hatching. In this study, we noticed a key developmental feature of saury development, that would attribute to the rapid growth of this species. In particular, *C. saira* and medaka exhibit several differences with respect to the timing of critical developmental events, including the onset of somitogenesis. Medaka, similarly to zebrafish, starts to form segmented somites after the completion of epiboly, the teleost version of gastrulation, and the appearance of primordial tailbud. Accordingly, developmental timetables of medaka and zebrafish are divided into ‘gastrula stages’ and ‘XX somite stages’, not overlapping with each other ^7,8,24^. In these species, somitogenesis starts and progresses in concert with the elongation of tailbud. In contrast, somitogenesis in *C. saira* begins in mid-epiboly stage, well before closure of the blastopore. Somite number increases along with the progress of epiboly. When the blastopore closes (100% epiboly), there are more than 10 somites at each lateral side of the embryonic body. At this point, tailbud is yet formed. It is also noteworthy that the Kuppfer’s vesicle, a small vacuole that has known to serve as an organizer of left-right asymmetry in organogenesis ^26,27^, appears at 50% epiboly and lasts until the blastopore closes. Presence of Kuppfer’s vesicle is no longer clear in the tailbud stages of *C. saira*, suggesting that molecular interactions underlying left-right asymmetry occurs mainly during epiboly stage in this species. Overall, these observations suggests that sauries exhibit ‘premature’ establishment of larval body structure that facilitates acute growth after hatching.

In the conventional view of vertebrate development, somitogenesis is supposed to occur after the completion of gastrulation. Gastrulation leads to the formation of mesodermal layer between the endoderm and the ectoderm, and after the closure of blastopore (future anal pore), the first somite pinches off at the most anterior edge of definitive presomitic paraxial mesoderm. As mentioned above, teleost models, such as zebrafish and medaka, are no exception – it is supposed that cells in the germ ring continue involuting movement in which the presumptive endoderm and mesoderm cell layers are formed underneath the ectodermal layer ^28^.

Remarkably, precautious somitogenesis has been reported in several teleost species, such as rainbow trout ^9^ and Midas cichlid fish ^25^. It is also noteworthy that recent studies using single-cell level imaging showed that, in zebrafish, endoderm and mesoderm arise from a single layer defined as mesendoderm that separates into two distinct germ layers after the completion of epiboly ^29^. This suggest that zebrafish mesoderm specification might not be finalized before the end of gastrulation. Together with these insights, our observation of *C. saira* suggest a variety in the relative timing of major morphogenetic events during early teleost embryogenesis.

## 4. Conclusions

In this study, we describe a first detailed description of embryonic development of a Belonid fish, the Pacific saury *C. saira*. The only additional fish studied so far in the same phylogenetic order is the medakas, and these two groups have diverged as early as 74 mya. Sauries and medakas, both of which have fully sequenced genome, have considerable differences in body size, habitats and behavior, and thus would serve as ideal models in the comparative studies of developmental genomics. We anticipate that developmental staging in this study will facilitate further progress of research on *C. saira*, which exhibits important features such as the nutritional contents, life cycles and behavior, which could have been impacted by the rapidly changing oceanic environment.

## 5. Experimental procedures

### 5.1 *C. saira* rearing conditions

Adults (originally from Maizuru, Kyoto, Japan) and most of the embryos of *C. saira* used in this study were sampled from the exhibition tank of Aquamarine Fukushima aquarium (Iwaki, Fukushima, Japan). Eggs were laid on the ropes that were hung inside the tank and inseminated immediately by males (Figure 1B)^2^. These ropes with fertilized eggs were removed from the exhibition tank and incubated in a running water tank in the backyard of the aquarium. These naturally spawned eggs were obtained at various developmental stages depending on the timing of fertilization. For microscopic observation, eggs were removed from the ropes and attaching filaments were trimmed with microscissors. Fertilized eggs and embryos were reared at 16 degrees in artificial sea water (Rei-sea Marine II, Iwaki Co., Ltd.) containing 50mg/L Streptomycin (Sigma-Aldrich).

To determine the developmental stages of cleavage and early epiboly, artificial fertilization was carried out by obtaining the eggs and sperm from mature adults. These gametes were mixed in a plastic container, followed by repeated washes with sea water to remove excess sperm.

For observation of late development, clutches of eggs were transported from Aquamarine Fukushima to Kansai University, Osaka. These eggs were placed in the petri dishes with artificial sea water containing Streptomycin and staged with reference to the approximate timing of developmental events observed in artificially fertilized eggs.

### 5.2 Microscopic observation of *C. saira* development

Early embryogenesis was observed and photographed using brightfield stereomicroscopes Zeiss SteREO Discovery.V12 (Zeiss) with a camera Axiocam 506 color (Zeiss) at Aquamarine Fukushima and stereomicroscope Leica M205C (Leica) with the digital camera Olympus DP74 (Olympus) at Kansai University. Hatched larvae were anesthetized with Ethyl *m*-aminobenzoate methanesulfonate (MS222, Fujifilm Wako Pure Co.) and mounted on the gel layer of 1% agarose in a petri dish filled with artificial sea water.

## Acknowledgments

We thank Rintaro Ishii, Toshiaki Mori and Hikari Yoshizato for their maintaining the Pacific saury colonies at Aquamarine Fukushima and their technical assistance for obtaining fertilized eggs. This study was supported by Grant-in-Aid for Scientific Research (C) (22K06243) to R.K. and by The NOVARTIS Foundation (Japan) for the Promotion of Science to R.K. and Mitsubishi Foundation to R.K.

